# Revisiting suppression of interspecies hybrid male lethality in *Caenorhabditis* nematodes

**DOI:** 10.1101/102053

**Authors:** Lauren E. Ryan, Eric S. Haag

**Author notes:** Correspondence: E.S. Haag, Dept. of Biology, Univ. of Maryland, 4094 Campus Dr., College Park, MD 20740.

## Abstract

Within the nematode genus *Caenorhabditis*, *C. briggsae* and *C. nigoni* are among the most closely related species known. They differ in sexual mode, with *C. nigoni* retaining the ancestral XO male-XX female outcrossing system, while *C. briggsae* females recently evolved self-fertility and an XX-biased sex ratio. Wild-type *C. briggsae* and *C. nigoni* can produce fertile hybrid XX female progeny, but XO progeny are either 100% inviable (when *C. briggsae* is the mother) or viable but sterile (when *C. nigoni* is the mother). A recent study provided evidence suggesting that loss of the *Cbr-him-8* meiotic regulator in *C. briggsae* hermaphrodites allowed them to produce viable and fertile hybrid XO male progeny when mated to *C. nigoni*. Because such males would be useful for a variety of genetic experiments, we sought to verify this result. Preliminary crosses with wild-type *C. briggsae* hermaphrodites occasionally produced fertile males, but they could not be confirmed to be interspecies hybrids. Using an RNA interference protocol that eliminates any possibility of self-progeny in *Cbr-him-8* hermaphrodites, we find sterile males bearing the *C. nigoni* X chromosome, but no fertile males bearing the *C. briggsae* X, as in wild-type crosses. Our results suggest that the apparent rescue of XO hybrid viability and fertility is due to incomplete purging of self-sperm prior to mating.

## Introduction

Interspecies hybrids can provide insight into the genetic mechanisms behind diversity of organisms, speciation, and the arisal of novel traits. Reproductive barriers limit gene flow between species and can be pre- or post-zygotic. Pre-zygotic barriers include behavioral isolation or gametic incompatibility. Post-zygotic reproductive barriers include hybrid lethality (at any pre-reproductive developmental stage) or sterility. It is this latter barrier that is the focus here. The Bateson-Dobzhanzky-Muller model proposes a genetic basis for hybrid incompatibility, whereby incompatibilities between heterospecific loci result in impaired function or non-function (DOBZHANSKY 1936). In addition, interspecies hybrids often manifest Haldane’s rule, where the heterogametic sex is more severely impacted, presumably because of sex chromosome hemizygosity (ORR AND TURELLI 2001). Darwin’s Corollary to Haldane’s rule is also observed in many cases, where there is asymmetric male viability in reciprocal crosses (TURELLI AND MOYLE 2007).

*Caenorhabditis* nematodes are an excellent system to study the genetic basis of reproductive diversity. The genus contains both gonochoristic (male/female) and androdioecious (male/hermaphrodite) species making it possible to study the variation of reproductive mode. The essence of hermaphroditism is limited spermatogenesis in the context of the XX female ovary. How development of the bisexual germ line is regulated has been studied heavily in *C. elegans* (e.g. DONIACH 1986; GOODWIN *et al.* 1993; ELLIS AND KIMBLE 1995; FRANCIS *et al.* 1995; ZHANG *et al.* 1997; CHEN *et al.* 2000; CLIFFORD *et al.* 2000; LUITJENS *et al.* 2000) and, more recently, in *C. briggsae* (CHEN *et al.* 2001; HILL *et al.* 2006; GUO *et al.* 2009; HILL AND HAAG 2009; BEADELL *et al.* 2011; LIU *et al.* 2012; CHEN *et al.* 2014). These two selfing species, while superficially similar, evolved self-fertility independently (KIONTKE *et al.* 2011) and achieve via distinct modifications of the global sex determination pathway (HILL *et al.* 2006; GUO *et al.* 2009; HILL AND HAAG 2009; CHEN *et al.* 2014).

The first studies comparing germline sex determination in hermaphrodite and female species used candidate gene approaches (HAAG AND KIMBLE 2000; HAAG *et al.* 2002; HILL *et al.* 2006; HILL AND HAAG 2009; BEADELL *et al.* 2011; LIU *et al.* 2012) and forward genetic screens (KELLEHER *et al.* 2008; WEI *et al.* 2014). More recently, however, the *C. briggsae* and *C. nigoni* system has opened the tantalizing possibility of using hybrids between them to identify factors distinguishing hermaphrodite and female germline sex determination (WOODRUFF *et al.* 2010; KOZLOWSKA *et al.* 2012). However, these efforts have been thwarted by extensive genetic incompatibilities. *C. briggsae X C. nigoni* hybrids are subject to both Haldane’s rule and Darwin’s Corollary to Haldane’s rule (WOODRUFF *et al.* 2010). Specifically, no viable male F1 are found when wild-type *C. briggsae* hermaphrodites are mated with *C. nigoni* males, but viable yet sterile males are produced when *C. nigoni* females are crossed with *C. briggsae* males. Surprisingly, after laying a few hybrid progeny, *C. briggsae* hermaphrodites mated with *C.nigoni* males are sterilized by the aggressive *C. nigoni* sperm (TING *et al.* 2014)

F1 females from both possible *C. briggsae X C. nigoni* crosses produce viable progeny only when backcrossed to *C. nigoni* (Woodruff et al. 2010). This has allowed introgression of marked *C. briggsae* chromosomal segments into *C. nigoni* (YAN *et al.* 2012; BI *et al.* 2015). These segments remain large in spite in backcrosses, and harbor a number of inviability and sterility loci, some of which impact germline small RNA pathways (LI *et al.* 2016). In addition, polymorphisms within *C. briggsae* and *C. nigoni* can impact the severity of F1 hybrid phenotypes (KOZLOWSKA *et al.* 2012).

As for its ortholog in *C. elegans* (PHILLIPS *et al.* 2005), the *C. briggsae him-8* gene is required for faithful segregation of the X chromosome during meiosis (WEI *et al.* 2014). This, in turn, greatly elevates the spontaneous production of XO self-progeny, which are male. Thus, while an unmated *C. briggsae* hermaphrodite will produce less than 1% males naturally, *Cbr-him-8* mutants produce approximately 15% males (WEI *et al.* 2014). Recently, Ragavapuram et al. (2016) reported that loss of *Cbr*-*him-8* function can rescue hybrid male lethality and lead to fertile F1 male progeny. As such males might allow F1 X F1 intercrosses and new backcross types for *C. briggsae* and *C. nigoni* hybrids, we sought to verify and extend these results. Surprisingly, using the same strains as Ragavapuram et al. (2016) but with methods that eliminate the possibility of males arising from selfing, we find no evidence of hybrid male rescue.

## Methods

### Strains

*C. briggsae* PB192 (*Cbr-him-8(vI88) 1;stls 20120 [Cbr-myo-2p::GFP;unc-119(=)]X*) was provided by Scott Baird. *C. nigoni* JU1422 (inbred derivative of wild isolate JU1375) was provided by Marie-Anne Félix. *C. nigoni* wild isolate EG5268 was the gift of Michael Ailion.

All strains were maintained on 2.2% NGM agar (WOOD 1988) with OP50 *E. coli* bacteria as food source. Strains are available from the Caenorhabditis Genetics Center.

### Cbr-fog-3 RNA interference

*Cbr-fog-3* template was made with the polymerase chain reaction (PCR) using NEB Taq polymerase with recommended concentrations of dNTPs and primers. Primer sequences (including the underlined T7 promoter required for subsequent *in vitro* transcription) are: Forward: 5’-TAATACGACTCACTATAGGGAGCCGACGAAGTTCT TGAAA-3’ Reverse: 5’-TAATACGACTCACTATAGGGCCCACCATGGTCTGCAGATC-3’. The PCR product was purified using a Qiagen QIAquick PCR purification kit. 780 ng of *Cbr-fog-3* PCR product was used as template in a T7 in vitro transcription reaction (Thermo Fisher Scientific) following the provided protocol. The resulting dsRNA was purified by ammonium acetate and ethanol precipitation. PB192 hermaphrodites were picked onto a separate plate at the L4 stage 12 hours prior to injection. 40 adult worms were mounted on agar pads and injected with *Cbr-fog-3* dsRNA at a concentration of 3000 ng/µL. The injected animals were moved to individual NGM plates seeded with OP50 *E.coli* plates after 8 hours of recovery time.

### Crosses

For interspecies crosses not employing *Cbr-fog-3(RNAi)*, PB192 hermaphrodites were purged, similar to the method of Ragavapuram et al. (2016).

*C. briggsae* mothers who could not self-fertilize were produced through RNAi targeting of *Cbr-fog-3* (CHEN *et al.* 2001). Progeny of injected PB192 mothers that appeared to have the feminization of germline (Fog) stacking phenotype after reaching adulthood were moved in small groups to new agar plates seeded with OP50 *E. coli* and allowed to sit for six hours to ensure that no self-progeny were produced. *C. nigoni* EG5268 males were added at a 2:1 ratio of males:hermaphrodites to each of the Fog plates and allowed to mate for 4 hours. All plugged animals were moved to individual plates. *Cbr-fog-3(RNAi)* PB192 were also crossed to *C. briggsae* AF16 males to test *fog-3* effect on fecundity.

### Microscopy

Routine maintenance and crosses were performed using a Leica MZ125. Analysis of male germ line morphology was done using differential interference contrast (DIC) optics on a Zeiss Axioskop 2 plus at 400x magnification.

## Results

We performed preliminary experiments with *C. briggsae him-8; myo-2::GFP X* hermaphrodites (strain PB192) that had been ostensibly purged of self-sperm by serial transfer over several days until embryo laying stopped, as done by Ragavapuram et al. (2016). The expectation was that roughly 15% of progeny from mating such purged hermaphrodites with *C. nigoni* males would be fertile males. Using the *C. nigoni* strain JU1422 in three different trials with 6-7 mothers each, 0/47, 3/50, and 0/116 progeny were male (1.4%). Because Ragavapuram (2016) used *C. nigoni* males of the African EG5268 strain, we next considered that the possibility that the unexpectedly infrequent males obtained above was a strain effect. Using the purging approach, 7/66 progeny were male in the first experiment with EG5268 males (11%). Of these, two were extremely small and infertile, similar to those observed when F1 males have a *C. nigoni* X chromosome (WOODRUFF *et al.* 2010). Because *him-8* mothers produce nullo-Xoocytes at an appreciable frequency, this is expected (RAGAVAPURAM *et al.* 2016). Five others were GFP+, robust, and fertile, and thus candidates for rescued F1 males. However, crosses with these putative F1 males indicated they were *him-8* homozygotes, and thus likely to be selfed offspring of *C. briggsae* PB192 mothers, perhaps derived from residual self-sperm that were resistant to purging (data not shown). A second attempt to generate hybrid males with EG5268 sires produced three males, all GFP- and small.

The above preliminary crosses produced fertile males at a frequency lower than the expected 15%. They also indicated that complete purging in *C. briggsae* may be more difficult to achieve than had been previously appreciated. To ensure that all progeny being scored were interspecies hybrids, and to allow use of younger, healthier mothers, self-sperm were ablated in PB192 by *Cbr-fog-3(RNAi)* via maternal injection (CHEN *et al.* 2001). From over 2000 hybrid F1 embryos laid, 16 viable male adults were obtained (Table 1). These males had fully formed tails and exhibited mating behavior, but were GFP- and unusually small. Attempts to backcross them to their siblings failed to produce any embryos. Consistent with this apparent sterility, all of these males lacked a fully formed germ line (often apparently completely absent).

**Table 1.** Phenotypes of progeny from *Cbr-him-8; myo-2:: GFP; Cbr-fog-3(RNAi)* mothers mated to *C. nigoni* EG5268 wild-type males.

| Total embryos | Total viable<br>hybrid progeny | Female progeny | Male progeny,<br>GFP(-) | Male progeny,<br>GFP(+) |
| --- | --- | --- | --- | --- |
| 2045 | 1065 (52.1%) | 1049 (51.3%) | 16 (0.8%) | 0 (0.0%) |

All of the above results are consistent with all of the F1 males obtained being derived from fertilization of a nullo-X *C. briggsae* oocyte by a *C. nigoni* male X-bearing sperm. This produces an X_*Cni*_O genotype known already to produce sterile males (WOODRUFF *et al.* 2010). They also indicate a lack of any viability or fertility of F1 X_*Cbr*_O males, contrary to the interpretation of Ragavapuram (2016). To be sure that the germline feminization of P0 *C. briggsae* hermaphrodites by RNAi did not suppress normal fertility in their sons, we verified that male offspring of *Cbr-fog-3(RNAi)* Fog mothers have normal fertility (data not shown).

## Discussion

In both *C. elegans* and *C. briggsae,* loss of *him-8* function specifically impairs X chromosome pairing (PHILLIPS *et al.* 2005; WEI *et al.* 2014), and unpaired *C. elegans* chromosomes are subject to meiotic silencing (BEAN *et al.* 2004). In addition, hemizygosity of the X chromosome is thought to underlie Haldane’s Rule in male-heterogametic systems. These observations led Ragavapuram et al. (2016) to hypothesize that *Cbr-him-8* mutant hermaphrodites produce X-bearing oocytes with altered X-linked gene expression that rescues X_*Cbr*_O F1 viability. As plausible as this mechanism is, we were unable to replicate the rescue of these males in our own hybrid crosses when all possibility of selfing is eliminated via *Cbr-fog-3(RNAi)*. The rare sterile males we do observe result from a *C. nigoni* X-bearing sperm fertilizing a nullo-X oocyte of the *C. briggsae* hermaphrodite, which occurs often in the *Cbr*-*him-8* PB192 strain. We therefore conclude that the previous report of X_*Cbr*_O hybrid male rescue was premature, and may be the result of incomplete purging of *C. briggsae* mothers. Perhaps importantly, the PB192 strain produces male self-progeny at a rate similar to that reported by Ragavapuram et al. (2016) as F1 hybrids.

We see some evidence for cryptic retention of a small number of self-sperm in putatively purged *C. briggsae* hermaphrodites. This may occur if sperm-derived major sperm protein (MSP)-mediated signaling is insufficient to stimulate ovulation (MILLER *et al.* 2001). *C. briggsae* nullo-X sperm are preferentially used last by hermaphrodites (LAMUNYON AND WARD 1997), and *Cbr-him-8* hermaphrodites produce an unusual number of them (WEI *et al.* 2014). Residual conspecific self sperm may therefore be highly enriched in those lacking an X, such that even a small number of them would tend to produce pure *C. briggsae* PB192 males that are both fertile and GFP+. This could contribute to the appearance of hybrid male rescue.

It remains possible that under some conditions fertile *C. briggsae* x *C. nigoni* hybrid males may yet be produced. If so, it will be crucial to demonstrate their hybrid nature by genotyping assays. Sufficient genome data now exist for *C. briggsae* (ROSS et al. 2011) and *C. nigoni* (LI *et al.* 2016) to make such assays, such as simple PCR of indel polymorphisms (KOBOLDT *et al.* 2010), rapid and inexpensive.

## Acknowledgements

We thank Scott Baird for the PB192 strain and for numerous useful discussions. This work was supported by NSF grant IOS-1355119 to ESH and by assistantships from the University of Maryland Biological Sciences Program to LER.

